# Stable isotope informed genome-resolved metagenomics uncovers potential trophic interactions in rhizosphere soil

**DOI:** 10.1101/2020.08.21.262063

**Authors:** Evan P. Starr, Shengjing Shi, Steven J. Blazewicz, Benjamin J. Koch, Alexander J. Probst, Bruce A. Hungate, Jennifer Pett-Ridge, Mary K. Firestone, Jillian F. Banfield

## Abstract

The functioning, health, and productivity of soil is intimately tied to a complex network of interactions, particularly in plant root-associated rhizosphere soil. We conducted a stable isotope-informed, genome-resolved metagenomic study to trace carbon from *Avena fatua* grown in a ^13^CO_2_ atmosphere into soil. We collected paired rhizosphere and non-rhizosphere soil at six and nine weeks of plant growth and extracted DNA that was then separated by density using gradient centrifugation. Thirty-two fractions from each sample were grouped by density, sequenced, assembled, and binned to generate 55 unique microbial genomes that were >70% complete. The complete 18S rRNA sequences of several micro-eukaryotic bacterivores and fungi were enriched in ^13^C. We generated several circularized bacteriophage (phage) genomes, some of which were the most labelled entities in the rhizosphere. CRISPR locus targeting connected one of these phage to a Burkholderiales host predicted to be a plant pathogen. Another highly labeled phage is predicted to replicate in a *Catenulispora sp*., a possible plant growth-promoting bacterium. We searched the genomes for traits known to be used in interactions involving bacteria, micro-eukaryotes and plant roots and found that heavily isotopically-labeled bacteria have the ability to modulate plant signaling hormones, possess numerous plant pathogenicity factors, and produce toxins targeting micro-eukaryotes. Overall, ^13^C stable isotope-informed genome-resolved metagenomics revealed that very active bacteria often have the potential for strong interactions with plants and directly established that phage can be important agents of turnover of plant-derived carbon in soil.

## Introduction

Plant-derived carbon provides the energetic basis for an intricate web of life in soil, among the world’s most complex microbial ecosystems. Heterotrophic soil organisms are sustained primarily by the carbon that is fixed by plants, released into the rhizosphere, and eventually flows into the surrounding soil, feeding other organisms or becoming soil organic matter.

The soil ecosystem is characterized by interactions between organisms across trophic levels, which may direct the fate of plant-derived carbon in soil. These interactions have been difficult to investigate because of the tremendous physical and chemical heterogeneity of soil and resulting vast biological diversity. Much of the recent work on soil microbiology has been sequence-based and focused on generating inventories of bacteria and archaea using 16S rRNA gene fragments (1, 2) or fungi using Internal Transcribed Spacer region sequencing (3, 4). Since DNA extracted from soil includes genes from virtually all organisms present, it is possible to use soil shotgun metagenome sequencing to profile soil communities, potentially with genomic resolution. This is important because genomes provide not only phylogenetic information but also a cache of functional predictions. However, relatively few studies have achieved genomic resolution of soil, largely due to enormous strain complexity and relatively even abundance levels (5–9). Bacterial genomes encode a huge variety of known and unknown genes, including those that comprise biosynthetic gene clusters (BGC). Such biosynthetic pathways hold technological relevance but are also fundamental for soil ecology, as their products could mediate inter-organismal interactions, including offensive interactions via antibiotics, inter-organism and mineral-interactions via siderophores, and signaling compounds (10, 11). While there has been a surge of interest in viral diversity and ecology, there have been relatively few studies documenting bacterial-phage interactions in soil (12–14). Also present in soil are fungi, protists and larger organisms that have been documented via 18S rRNA gene sequencing (15, 16) and more classical methods (17, 18). While some studies have documented bipartite interactions, such as fungi-bacteria (19, 20), bacteria-phage (12, 21), plant-fungi (22, 23), detailing complex multi-trophic interactions in soil remains a huge challenge.

Lacking from most studies of soil to date is information about how nutrients flow among soil community members. Stable isotope probing (SIP) has been used for decades to identify interactions and follow the flow of elements through communities. SIP studies have been conducted in a variety of ecosystems, including hot springs and the animal gut, using a range of isotopes and labelled substrates (24–27). SIP studies can utilize carbon fixing organisms to generate biomass and complex mixes of compounds to investigate general processes such as decomposition of litter or life in the rhizosphere (28–31). Stable isotopes can also be traced between trophic levels allowing the study of microbial predation and phage activity (32, 33). A recently developed variant, quantitative SIP (qSIP), makes these measurements possible at the individual or population genome scale. This is facilitated by comparing the taxon-specific density for each sequenced entity in the labelled and unlabeled fractions (34, 35). From this data we estimated the gross growth rate of organisms on labelled carbon substrates.

To better understand the soil inter-organismal interactions that are initiated by the input of plant-derived rhizodeposits, we combined stable isotope probing and genome resolved metagenomics. We grew common wild oat grass, *Avena fatua*, for six and nine weeks in a ^13^CO_2_ enriched atmosphere and tracked the isotopically enriched plant carbon as it was released to the surrounding soil community via exudation, decay of root biomass, and direct biotic transfer through pathogen attack. Over the course of our experiment, we expected the plant-derived carbon to move through trophic levels, including primary consumers, predators, parasites, saprotrophs, and their phage. By separating extracted DNA based on density we determined which organisms consumed the isotopically heavy plant-derived carbon and incorporated it into their genomes during replication. We used assembly and binning of genomes to analyze the encoded genes at the organismal level and identify traits that may reflect how organisms might interact with one another. We targeted interaction signatures such as genes for the production of plant hormones and modulation of hormone concentrations, secretion systems, and secondary metabolites. Assembled genomes enabled the prediction of whether an organism is a plant growth-promoting bacteria (PGPB) or a pathogen (36, 37).

## Results

### Labelling

*A. fatua* plants were grown in a continuously regenerated ^13^CO_2_ environment. One plant was harvested after six weeks and a second after nine weeks. Samples were collected from the rhizosphere (within 2mm of the root) as well as soil regions from which roots were excluded (see Methods); these are referred to as “bulk” samples. We also collected a sample of soil prior to planting. This sample is also denoted as “bulk.” After six weeks of growth, the plant shoots were highly labelled (∼ 94 atom% ^13^C) (38). DNA was extracted from both the rhizosphere and bulk soil samples. We compared the density separation of the rhizosphere DNA to the bulk soil DNA to identify un-enriched DNA (light), a mixture of enriched and unenriched DNA (middle), and highly ^13^C-enriched (heavy) then these fractions were shotgun sequenced (Supplemental figure 1 and Supplemental table 1). After assembly of the individual metagenomes, it appeared that the heavier fractions (middle and heavy) had larger assemblies and higher N50 values than the assemblies of the more complex light fractions (Supplemental table 1).

### Soil community

Given that it is currently impossible to bin genomes for all organisms in a soil sample, yet relatively extensive reconstruction of genome fragments is possible, we used an assembled marker gene approach to approximate microbial community composition. For this analysis we used ribosomal protein S3 (rpS3), which is in single copy on bacterial and archaeal genomes and has been used to profile microbial communities for phylogeny and abundance (39, 40). From each sample we identified the rpS3 and dereplicated the sequences to a level of 99% nucleic acid identity (41). The resulting 314 distinct rpS3 sequences represent bacteria from both well characterized and understudied clades (**Figure 1**).

**Figure 1.**
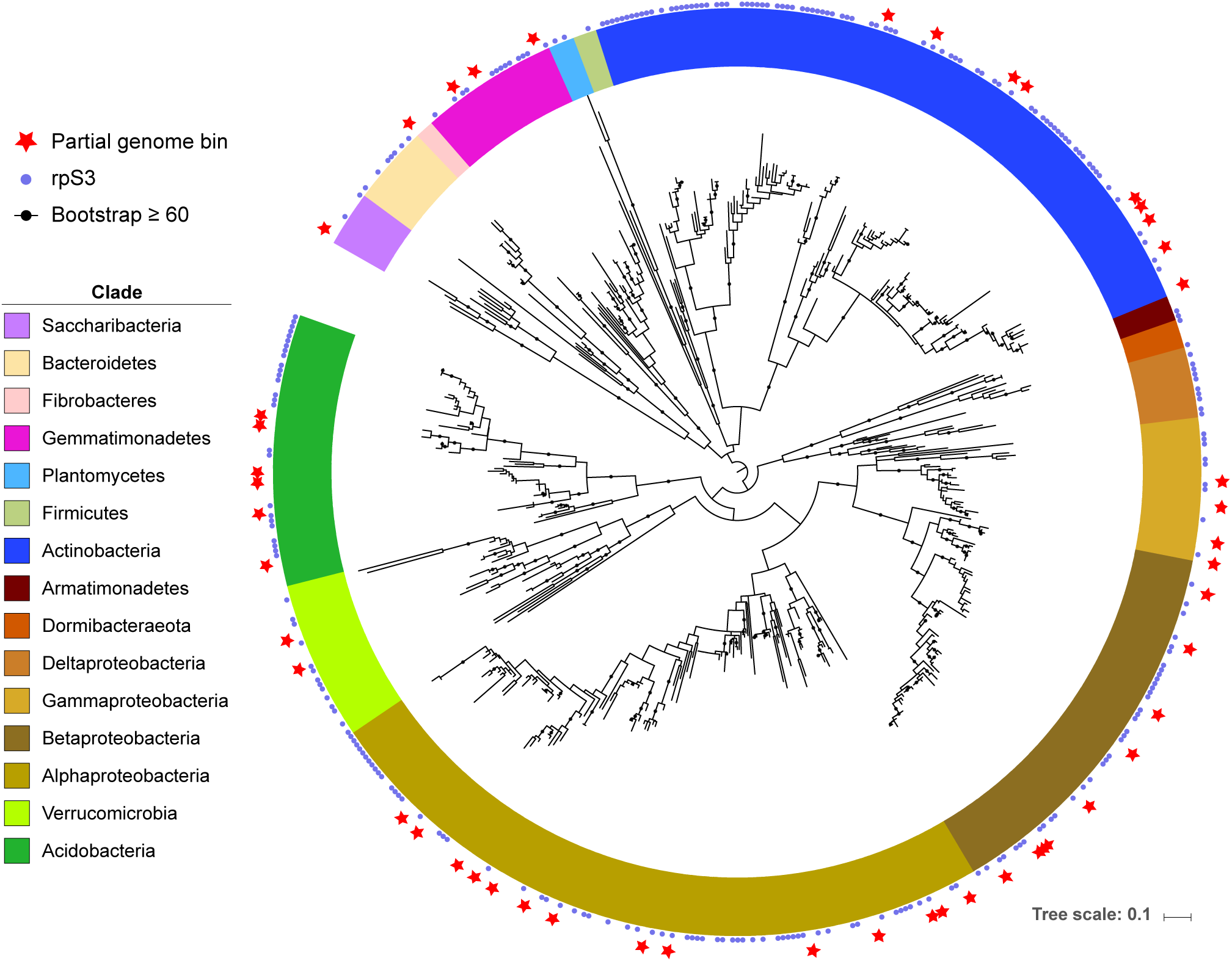
Phylogenetic tree of bacterial rpS3. Soil and rhizosphere-derived metagenomic bacterial bins with an rpS3 gene and unbinned scaffolds are marked. Bacterial clades are highlighted in different colors.

To study community dynamics across samples and fractions we mapped the reads from each sample and fraction to each scaffold containing the rpS3 gene. We used the coverage of the scaffolds as a proxy for the organism’s relative abundance. The fractions and samples show a visual community separation (**Figure 2**). The bulk samples separate into a light fraction (tan) cluster and a heavier fraction (brown) cluster. This is due to lower-GC organism’s DNA in the light fraction and the higher-GC DNA in the heavier fraction. The bulk light fraction and rhizosphere light fraction samples group together. The rhizosphere middle fraction separates from the light fractions (both rhizosphere light and bulk light fractions) and the bulk soil heavier fraction as it contains DNA from the high-GC organisms that did not incorporate the label and the lower-GC organisms that were labelled sufficiently to increase the density of their DNA.

**Figure 2.**
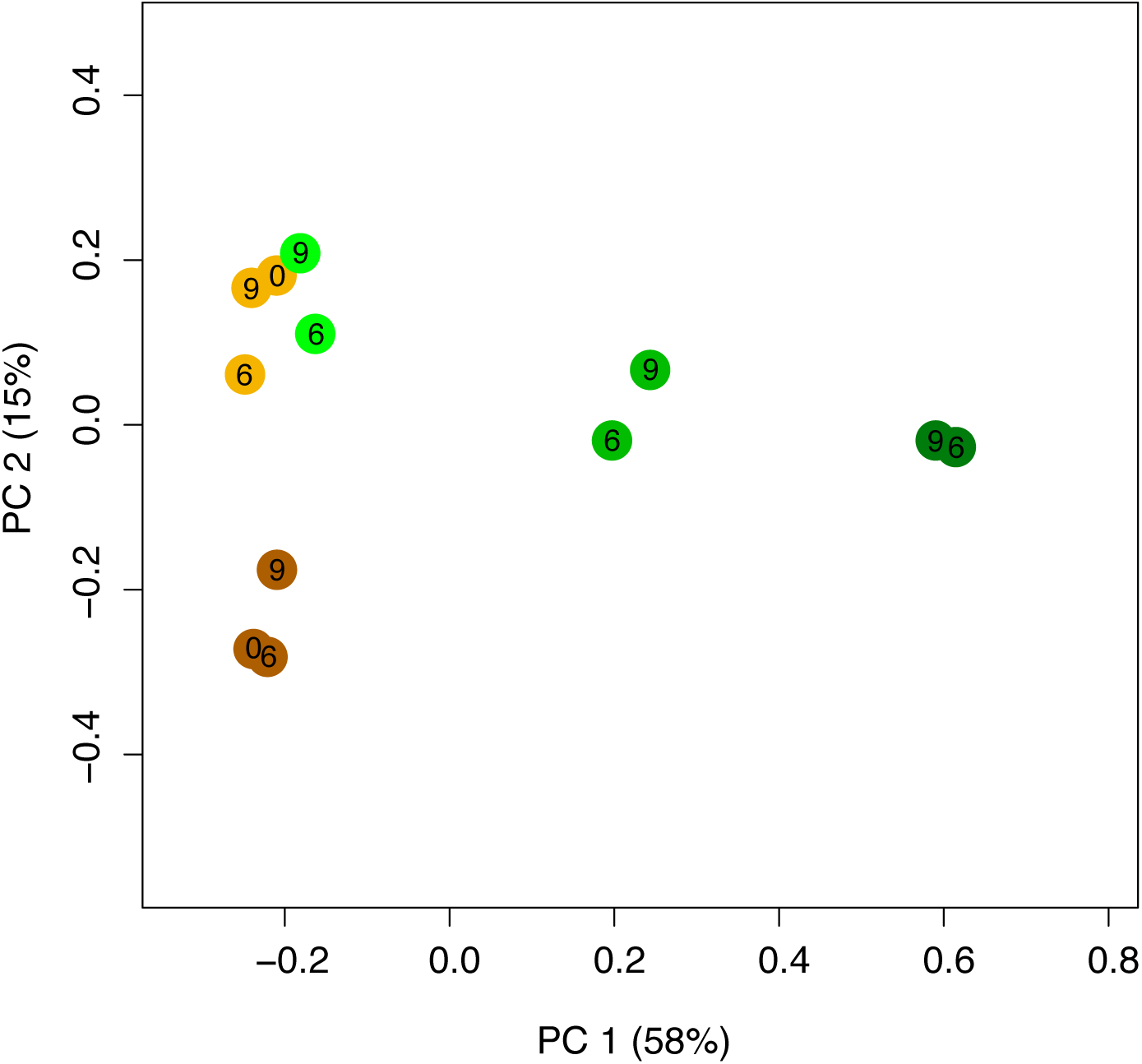
PCoA of bacterial rpS3 from different samples and SIP fractions. The brown dots represent bulk samples (the dark brown are the heavier fractions and the tan are light fractions). Green dots represent rhizosphere SIP fractions: the darker the green the heavier the sample. Numbers inside the dots correspond to the week of sampling.

Finally, the rhizosphere heavy fractions separate from all other fractions. The bacteria present in this sample are almost entirely high-GC organisms that incorporated ^13^C into their DNA.

From our 12 metagenomes, we reconstructed and binned 55 dereplicated, partial bacterial genomes that were ≥70% complete with ≤10% contamination, as measured by the inventory of 51 single copy genes (**Figure 3**; Supplemental figure 2).

**Figure 3.**
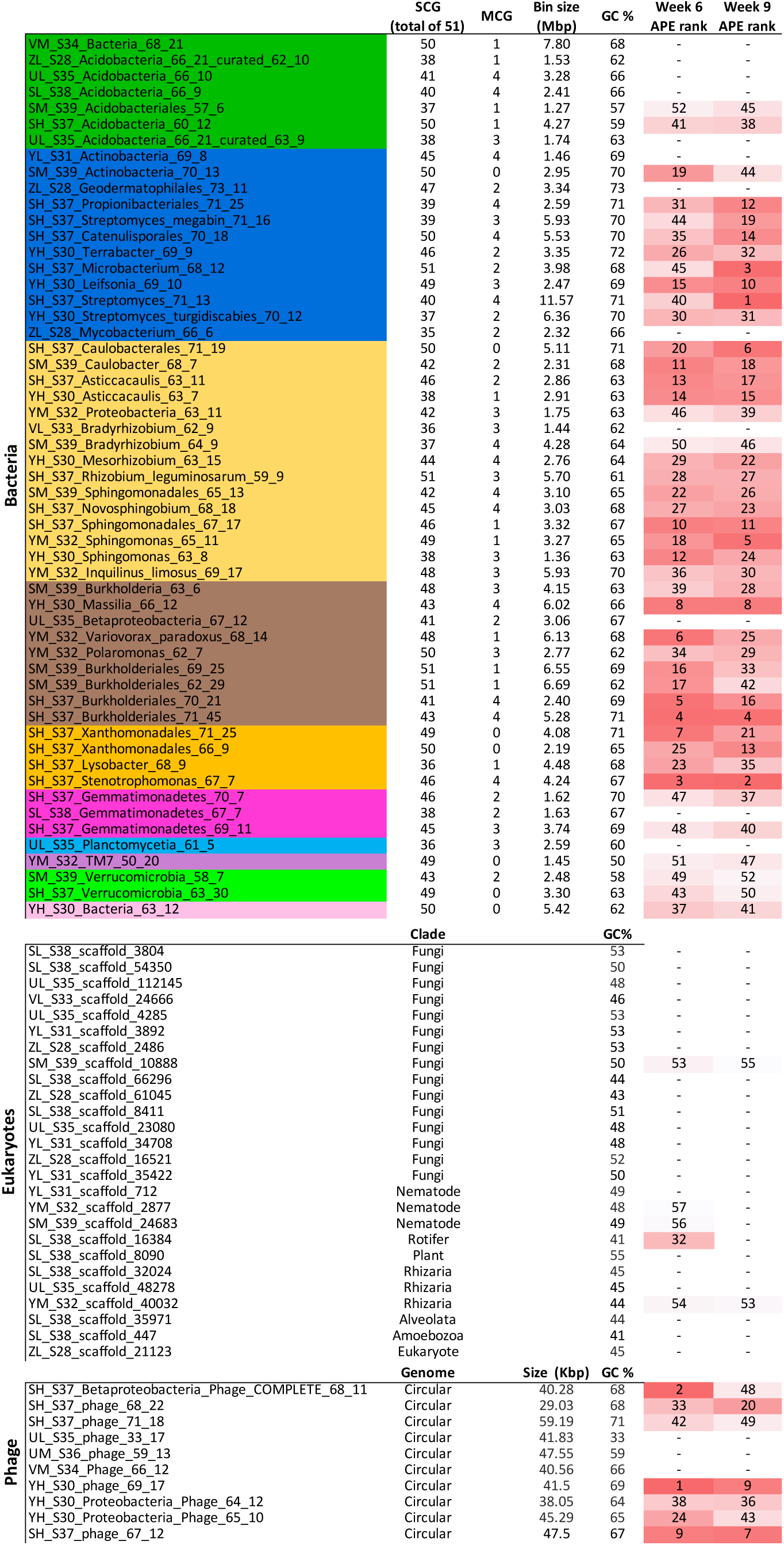
Genome and isotope labelling statistics. Genome statistics for the bacteria (colored by clade following the color scheme from Fig. 1), eukaryote scaffolds containing 18S rRNA genes, and complete phage genomes. The rank APE (highlighted with red heat maps for each column) is derived from the qSIP calculations, and is shown in ascending order for week 6 and 9. The bin completeness and contamination are presented as the number of 51 single copy genes (SCG) and number of multicopy genes (MCG).

In addition to bacteria, we detected a number of eukaryotes and phage. We reconstructed 27 complete 18S rRNA gene sequences from soil eukaryotes and used them for identification (**Figure 3**; Supplemental figure 3). Apart from the *A. fatua* and a single moss, the remaining 25 micro-eukaryotes fall into a variety of soil clades including Amoebozoa, Fungi, Metazoa (nematodes and rotifers), Rhizaria, and Alveolata. We also identified phage-derived DNA in our samples and reconstructed 10 complete, circularized phage genomes (**Figure 3**).

We used quantitative stable isotope probing (qSIP) to estimate ^13^C atom percent excess (APE) for each taxon (34, 35). The qSIP method relies on tracking the shift in density, calculated by coverage in the different fractions, of a genome between unlabeled (bulk) and labelled (rhizosphere) samples. We mapped the reads of all samples against our dereplicated suite of 55 genome bins. We used the coverage of the scaffolds containing the 18S rRNA as a proxy for eukaryotic genome coverage. Because of the lack of replicates and small number of fractions we chose a conservative cutoff of 2.5% APE (see Methods), any entities with an APE higher than this cutoff were interpreted as having detectably incorporated the ^13^C tracer (**Figure 4**). We report the rank in ascending order of APE for each sequence in **Figure 3**. Many of the phage we identified appear to be highly labelled. Bacteria were also highly labeled, and some of eukaryote sequences were labeled, although to a lesser degree than the bacterial genomes (**Figure 4**).

**Figure 4.**
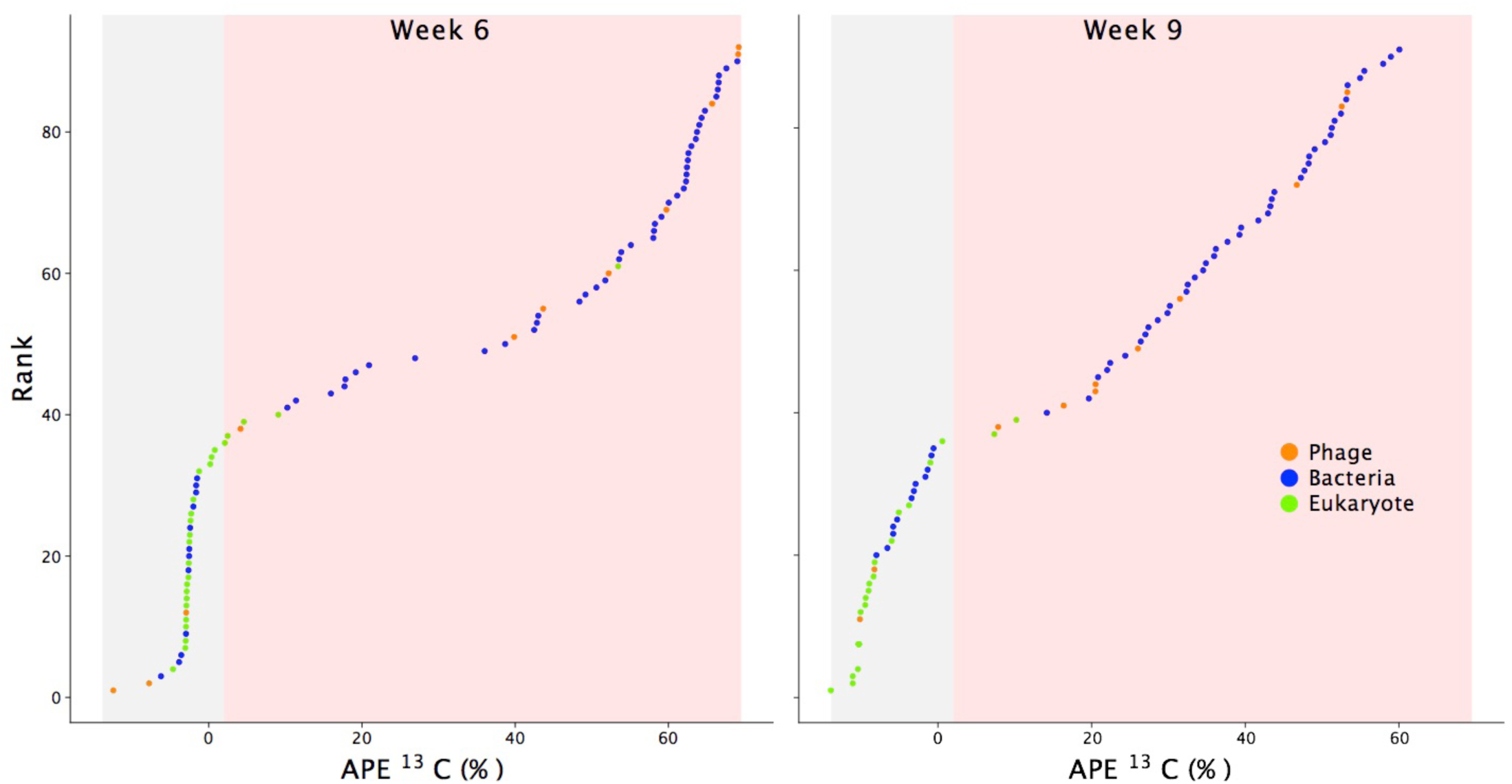
The rank of soil-derived phage genomes, bacterial genome bins, and scaffolds encoding eukaryotic 18S rRNA genes in week 6 and 9 in order of their atom percent excess (APE) based on the qSIP calculations. The gray region indicates unlabeled entities and the red indicated predicted labelled DNA, the labelling cutoff is explained in the methods section.

We calculated the gross growth rate for each taxon on labelled carbon. These taxon-specific growth rate estimates can be thought of as a measure of which individual organismal populations in the rhizosphere grew on plant-derived carbon. The gross growth rates indicated bacterial and phage growth but limited eukaryotic growth on plant-derived carbon. Indeed, only a few specific bacteria and phage had high gross growth rates on plant derived-carbon, and many taxa showed little to no growth (Supplemental figure 4).

We analyzed our genomes for a suite of genome sequences that could code for pathways involved in inter-organismal interactions. The possible machineries involved in these interactions provide the foundation for a conceptual understanding of the carbon flow and multitrophic interactions in these soil samples, as shown in **Figure 5**.

**Figure 5.**
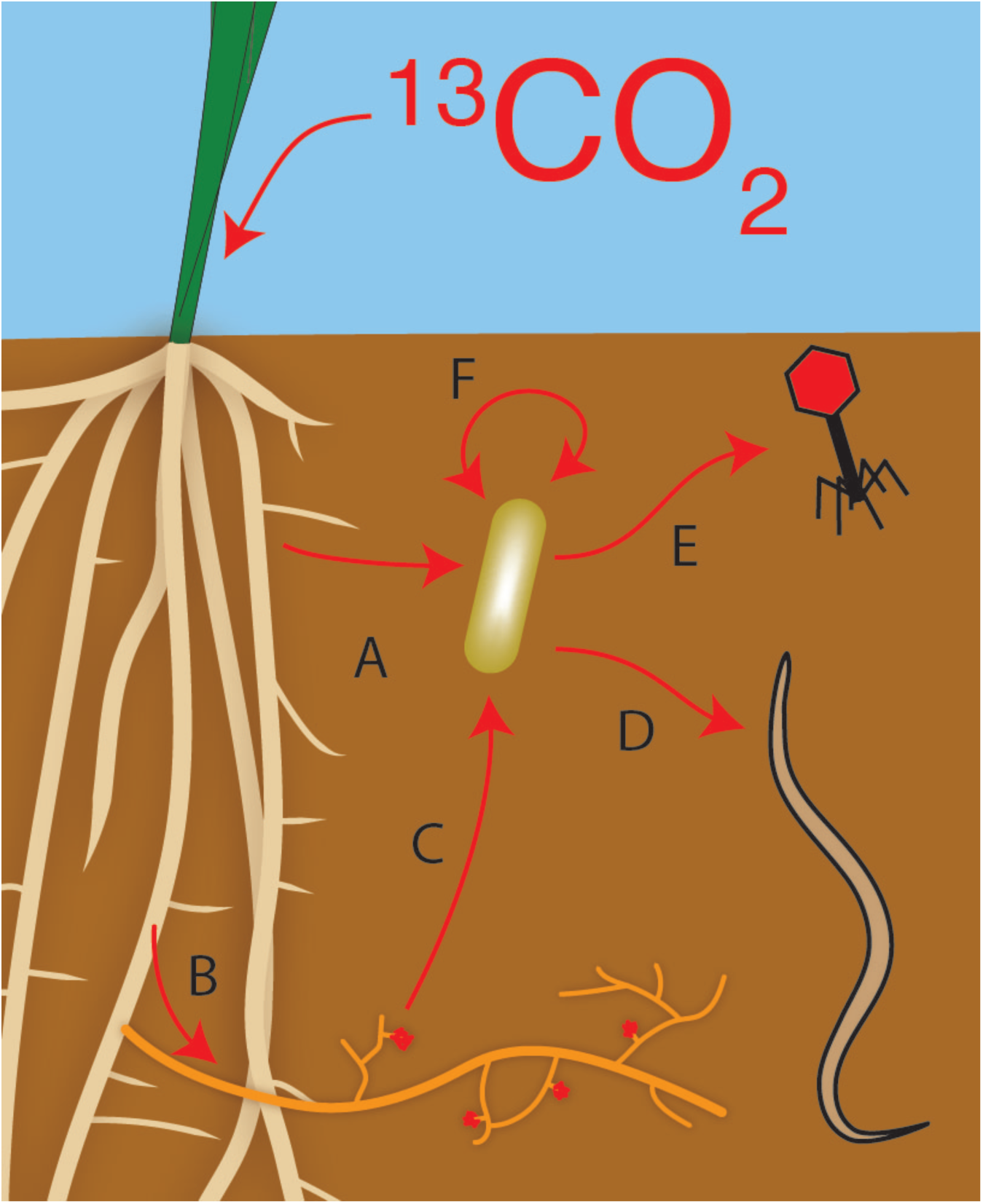
Conceptual diagram of carbon flow and community interactions in the rhizosphere. Red lines indicate possible flow of 13C label into and through DNA. Bacteria consume plant exudates or actively infect the plant to receive carbon (A). Fungi may also be infecting and/or decomposing dead plant material (B). Rhizosphere dwelling bacteria may be interacting with live fungi or breaking down dead fungal biomass (C). Bacterivorous micro-eukaryotes may have consumed rhizosphere bacteria. (D). Bacteriophage may have infected rhizosphere dwelling bacteria (E). Bacterial genomes were also rife with systems made for communication and competition between bacteria (F).

### Plant-soil community

Although plant-derived carbon is the main source of ^13^C used by the soil community, other organisms may be able to fix CO_2_. However, we found no evidence for carbon fixation pathways in the bacterial genomes and the lack of density shift in the bulk samples indicates that this effect was undetectable (Supplemental figure 1). Many of the bacterial genomes encoding plant interaction mediating genes and pathways were highly labelled, suggesting an intimate relationship between the plant roots and bacteria.

Bacteria that are closely associated with plants may degrade plant infection signaling hormones to avoid detection during plant colonization. Many of the bacterial genomes we binned encoded the ability to hydrolyze salicylic acid (**Figure 6**), a common phenolic plant hormone used in pathogen defense signaling (42). However, this protein can also be used to degrade other phenolic compounds (43). Thus, we examined the nearby genomic regions for clues about the function of the gene. In one instance, the salicylate hydroxylase gene in the Microbacterium_68_12 genome is surrounded by a variety of glycosyl hydrolases and esterases that act on plant cell wall polymers. Additionally, many of the rhizosphere-dwelling organisms encode the ability to degrade another pathogen defense hormone, nitric oxide gas (44).

**Figure 6.**
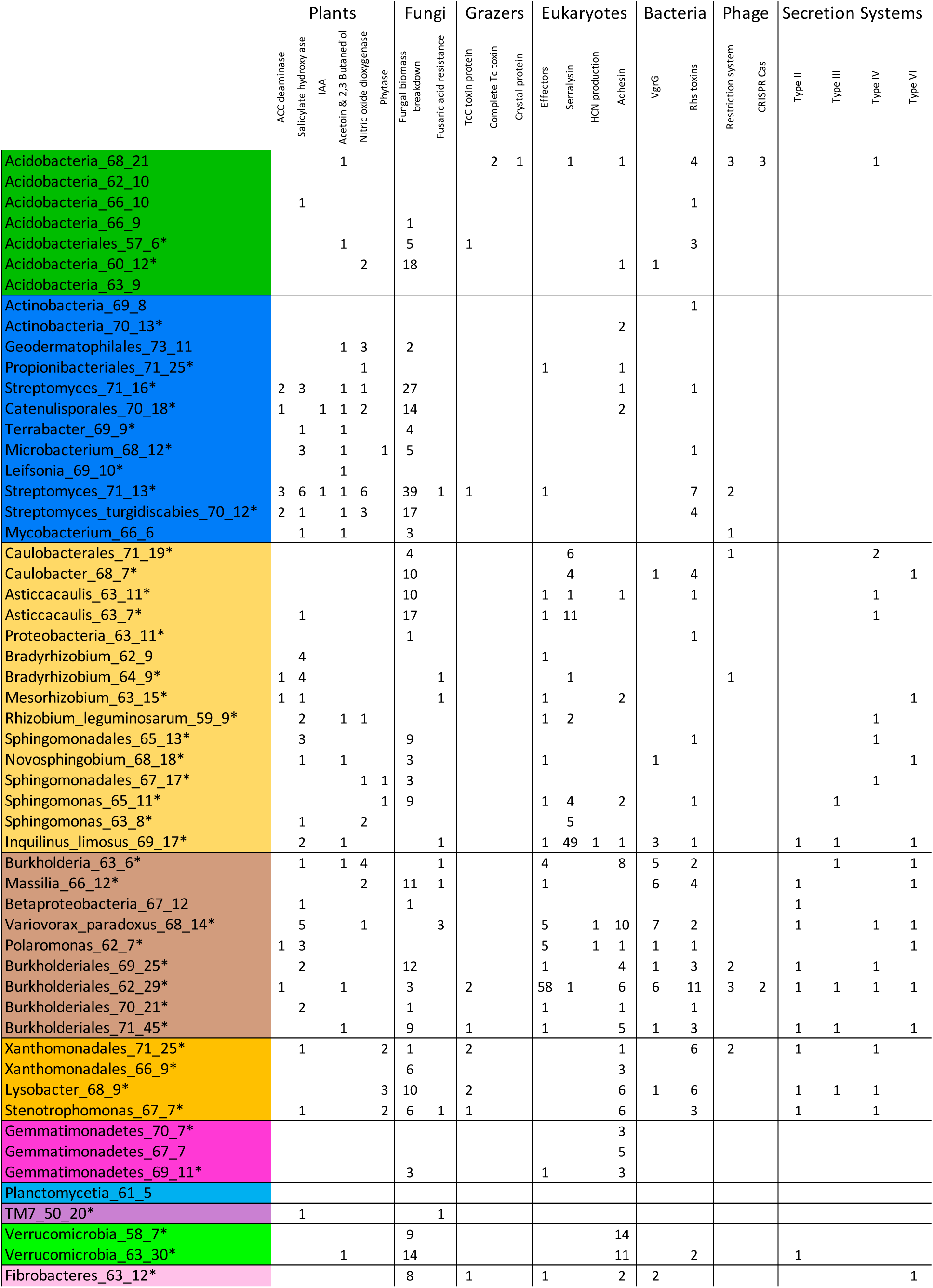
Possible inter-organismal interactions encoded on the bacterial genomes. Numbers indicate the number of individual genes or mostly complete pathways predicted to be used in inter-organismal interaction. Asterisks signify genome bins with detectable ^13^C label.

Some bacteria, especially PGPB, promote plant growth through the production of hormones and other compounds. Two of the bacterial genomes we reconstructed encoded the pathway for indole-3-acetic acid production, which increases plant growth and induces a variety of other physiological changes (**Figure 6**). Eight genomes encode 1-aminocyclopropane-1-carboxylate (ACC) deaminase (**Figure 6**), which prevents ACC from being converted to ethylene in the plant. However, ACC deaminase is also involved in the generation of propionate. In the Streptomyces_71_13 genome the ACC deaminase gene is surrounded by pectin lyases, pectinesterases, and a BGC. We identified 18 genomes with the pathways for the production of acetoin and/or 2,3-butanediol from pyruvate (**Figure 6**). These bacterially produced volatile organic compounds (VOCs) diffuse through soil and can act as growth-promoting factors and stimulate the plant systemic defense. Although these compounds increase resistance to plant pathogens, the pathway can also be involved in the anaerobic fermentation of glucose (45, 46).

Microbes can also promote plant growth through nutrient generation or mobilization. We did not identify any N_2_ fixing pathways in the genomes or on the unbinned scaffolds. Some PGPB are able to mobilize phosphorus for use by the plant, but these genes cannot be distinguished from central metabolism and common enzymes. However, microbially produced phytases release phosphorus from phytate, a phosphorus storage compound common in soil but inaccessible to mature plants (**Figure 6**) (47). Several of the genomes encoded biosynthetic pathways to produce siderophores (Supplemental figure 5). In addition to iron acquisition, siderophores can complex iron and other metals, thus promoting the release of phosphate from insoluble soil-associated minerals (48).

Some of the genomes appeared to encode secretion systems associated with bacteria-eukaryotic interactions. Six genomes encode multiple genes from type III secretion systems, which are known to be important in symbiotic colonization and infection of plants, fungi, single-celled eukaryotes, and animals (49). We do not know the intended targets of the type III secretion systems because of the diversity of possible hosts. However, we identified 58 type III effector proteins with sequence homology to known plant pathogen effector proteins (mostly from *Ralstonia, Pseudomonas* and *Xanthomonas*) encoded on the Burkholderiales_62_29 genome (**Figure 6**). We also identified 13 genomes with probable type IV secretion systems (**Figure 6**), which are used in conjugation and injection of proteins or DNA/protein complexes into eukaryotic cells (50).

### Bacteria-Microeukaryotes

Several of the bacterial genomes we assembled encode systems that may be intended for interacting with soil fungi. Ten of the partially complete genomes encoded fusaric acid resistance proteins, which protect from the antibiotic produced by *Fusarium* species (**Figure 6**). The Variovorax_paradoxus_68_14 genome appeared to encode three fusaric acid resistance modules; and, one was near an esterase and phospholipase C which hydrolyzes phosphatidylcholine, an important fungal phospholipid (51). In one Streptomyces_71_13 genome the fusaric acid resistance module is near two glycoside hydrolases which may act on fungal cell walls. Included in this region is a ceramidase, which hydrolyzes glucosylceramides, a necessary metabolite for *Fusarium* pathogenesis and morphology (52). Another fungal antibiotic that these bacteria may encounter is penicillin, possibly produced by the *Eurotiomycetes* detected in the soil. Many bacterial genomes encoded penicillin resistance genes. Another possible indication of bacterial-fungal relationships is bacterial genes for decomposition of fungal compounds. The Streptomyces_71_13 genome encodes 39 enzymes that target fungal biomass, 30 of which have an identifiable secretion signal indicating they are exported extracellularly (**Figure 6**).

There is some evidence for bacterially produced defenses against grazing. The majority of work on bacterial grazing defense has focused on nematodes, but we assume similar strategies could be effective against other micro-eukaryotes. Bacteria often use extracellular polymeric substance (EPS) production, specific secondary metabolites, and active infection to deter grazing (53). EPS production is common in soil bacteria, but difficult to infer from genome information. One indication of EPS formation is the production of proteinaceous adhesins (54). Many of the genomes we investigated encoded adhesins, with two Verrucomicrobia (Verrucomicrobia_58_7 and Verrucomicrobia_63_30) and the Variovorax_paradoxus_68_14 genomes encoding the most (**Figure 6**). Several genomes encode the pathway to produce hydrogen cyanide (HCN; **Figure 6**). Eight of the genomes encode a insecticidal toxin subunit TcC which is lethal to certain insects, and possibly nematodes, on its own (**Figure 6**) (55). However, the Acidobacteria_68_21 genome encoded two complete Tc insecticidal toxin modules. In addition, in the same genome, we identified an insecticidal crystal protein related to the bt toxin from *Bacillus thuringiensis* (**Figure 6**).

### Bacteria-Bacteria

In addition to bacterial interaction, cooperation, and competition with eukaryotes there may be indications of inter-bacterial connections in soil. Some of the best characterized mediators of inter-organismal interactions are signaling molecules including acyl-homoserine lactones, autoinducing peptides, indoles, gamma-butyrolactones, and a variety of other compounds (Supplemental figure 5). In addition, many genomes encode one or more quorum quenching genes, which may act to either degrade self-produced quorum molecules or as a means to disrupt other bacterial species communications.

We identified a large number of biosynthetic gene clusters in the bacterial genomes, especially polyketide and nonribosomal peptide biosynthetic gene clusters (Supplemental figure 5). Several *Streptomyces sp*. encoded many BGCs and several Burkholderiales and Acidobacteria genomes also encoded a high number of secondary metabolite clusters, though this is less surprising in light of recent findings in other soils (11, 56). Some of the BGCs were related to known clusters, such as those that produce the broad spectrum antibiotic bicornutin from *Xenorhabdus budapestensis* and the sesquiterpene antibiotic albaflavenone, but many were novel. Some of the BGCs were located near other genes of interest for instance, the Streptomyces_turgidiscabies_70_12 genome encoded three BGCs near ten plant cell wall hydrolysis proteins. Evidence for competition between closely related strains comes in the form of bacteriocins, antibiotics that act on closely related bacteria. These were found in nearly 40% of the genomes. Ten genomes encoded type VI secretion systems, which inject effectors into neighboring bacteria (**Figure 6**). Many genomes had multiple VgrG proteins, the tip of the needle and effector transporter, near large proteins with Rhs repeat domains which may function as bacterial toxins (**Figure 6**) (57). The Rhs proteins were not limited to the bacteria with type VI secretion systems and may be exported in other ways.

### Bacteria-Phage

Many of the phage we identified appeared to be highly labelled (**Figure 4**). In the week 6 sample the two most labelled entities were phage (**Figure 3**). We focused on circularized phage genomes (not integrated into the host genome as a prophage where they could become ^13^C labelled through host growth alone) as these are likely complete genomes and the product of active infection during our experiment.

We identified Burkholderiales_62_29 as the possible host for one of the most highly labelled phage, Burholderiales phage 68_11, based on the match between a CRISPR-Cas spacer and the complete phage genome. The spacer hit the large terminase subunit with two mismatches (33 bp total). It appears the phage may be capable of a lysogenic life cycle because of the presence of a serine recombinase and a possible induction region consisting of a histone-like protein H-NS and an AT rich region flanking a lambda repressor-like gene.

The other phage-host connection we identified was based on a possible recent lateral gene transfer event. The phage Catenulispora phage 69_17 was highly labelled, and ranked as the 1^st^ and 9^th^ most labelled entity in week 6 and 9 respectively (**Figure 3**). The probable host, Catenulisporales_70_18, encodes a glycoside hydrolase (GH25) which is very similar to one found in the complete phage genome, possibly indicating the phage may have acquired the gene from this bacterial population. The two proteins share 77% amino acid identity across the protein and a phylogenetic tree of the proteins indicates that the phage and bacterial proteins are more related to one another than to other publicly available bacterial sequences (Supplemental figure 6). Homologs exist in many *Catenulispora* genomes but not in closely related phage genomes, consistent with recent acquisition of the gene by lateral transfer. The acquired gene may be a lysis factor, as GH25 breaks down peptidoglycan. The phage may have a lysogenic life cycle based on the presence of a gene annotated as a tyrosine recombinase and induction regulation genes, a lambda repressor-like gene and a gene containing a GntR family transcriptional regulator domain and an UTRA domain which regulates the activity of transcription factors in response to small molecules (58, 59).

The remaining eight complete phage genomes cannot be linked to a specific host. Many encode DNA methylation genes that may protect the phage DNA from detection or destruction by host antiviral systems. Some phage have additional genes encoding complete restriction modification systems. Several phage genomes encode the ability to manipulate the composition of the bacterial nucleotide pool, including one phage that encodes for the synthesis of 5-hydroxymethyldeoxyuridylate, a special nucleotide that replaces thymine in the phage genome to evade antiviral defense systems. Another phage may encode auxiliary genes, proteins related to those involved in glycosylation and biosynthesis of phosphatidylinositol mannosides. Of the 55 draft bacterial genomes, only two contained identifiable CRISPR-Cas systems, the Burholderiales described above and Acidobacteria_68_21, which encoded two Cas type III-B systems and a type I-C system (**Figure 6**).

## Discussion

By combining stable isotope probing with genome resolved metagenomics, we identified a number of possible interactions among soil dwelling organisms; these possible interactions become interesting targets for further exploration and testing to determine if they are indeed operating and important in the soil/rhizosphere. The process of density separation yielded better assemblies which likely improved binning by providing more accurate tetranucleotide frequencies and coverage values. Through the generation of genome bins we were able to provide clues regarding the ecological role of these organisms and to develop ideas about the movement of carbon through the system.

A better understanding of holistic rhizosphere ecology may help us develop better agricultural techniques and biotechnological products. Some of the bacteria in our system may be acting as PGPB, through their ability to release nutrients, help in water stressed conditions, or by repelling plant pathogens. By understanding the different trophic networks, we may begin to develop multi-domain plant growth-promoting consortia that could include phage of plant pathogens, fungi, and microeukaryotes. From only 55 bacterial genomes we detected 400 BGCs, many of these compounds could be novel antibiotics, antifungals, insecticides, herbicides, or pharmaceuticals.

Based on ordination of rpS3 genes from different samples and isotopically enriched fractions, it appears that the supply of labeled plant root carbon to soil identifies organisms from the background soil community forming visually distinct assemblages of labelled rhizosphere organisms. Also, within the rhizosphere, a large portion of organisms were not directly responsive to the influx of plant-derived carbon and their communities were indistinguishable from bulk samples. Over time, successional shifts may have occurred, but the resolution of this method is not sufficient to detect them. The traditional 16S rRNA gene tag sequencing could identify effects of sampling time point on microbial composition in a parallel experiment (60), but this approach is unable to provide genomic insights. In addition to bacteria, we identified a number of soil eukaryotes, and though this number does not scratch the surface of soil eukaryotic diversity, the assembled metagenomes provide complete 18S rRNA sequences without the primer bias inherent with tag-based methods. We identified 10 complete phage genomes which likely represent some of the most abundant phage in our system. This is a small number compared to the total diversity of phage likely present in the soil, yet the complete genomes allow us to predict lysogenic lifestyles and to identify possible hosts of the dominant phage.

By assuming linear growth over the course of the experiment, we can not only estimate enrichment, but also derive gross growth rate estimates for prokaryotes, micro-eukaryotes, and phage that rely on plant-derived carbon. The uncertainties of these estimated rates mirror the uncertainties in measuring the abundance of metagenome-assembled genomes in soil. The use of larger numbers of density fractions results in much reduced uncertainties in growth rate estimates (61). However, as expected, both phage and bacteria showed higher average growth rates than the slower-growing eukaryotes. These quantitative metrics enable linking previously undescribed populations of rhizosphere microbiota to plant-derived C-fluxes in an intact plant-soil environment. Future developments in this technique will allow us to better understand complex carbon utilization networks in soil.

Many of the most highly labelled organisms were those with suspected plant interaction systems and may span the spectrum from mutualist to pathogen. By analyzing genomes with >70% completeness we were able to not only look for genes or pathways involved in interaction but also investigate the genomic context for additional information to help predict the purpose of these genes (62, 63). For instance, in the Streptomyces_71_13 genome the ACC deaminase gene is adjacent to plant cell wall hydrolysis genes and the biosynthetic pathway for producing ectoine, a compatible solute common in PGPB (40). This region of the genome may allow the limiting of a stress response in the plant by lowering ethylene levels and producing an osmoprotectant which would ensure greater plant growth (64, 65). The plant cell wall digestion enzymes may enable this PGPB to invade the plant to form a closer mutualistic association with the roots. Apart from that specific region of the genome, Streptomyces_71_13 also encodes a number of other pathways and genes known to be important in PGPB including the production of indole-3-acetic acid and 2,3-butanediol. The possible close association with plant growth promotion may explain why Streptomyces_71_13 was the most labelled entity in week 9.

In addition to possible PGPB we also identified probable plant pathogens. Based on the presence of a type III secretion system and plant effector proteins, Burkholderiales_62_29 may act as a plant pathogen, which may have enabled the assimilation of large amounts of plant-derived carbon making this genome the 17^th^ most labelled entity in week 6. To further support this hypothesis, we referred to the transcription of the effector proteins from a related study of the *Avena* rhizosphere metatranscriptome which used the Burkholderiales_62_29 genome as a reference, eight of the 58 effectors were statistically upregulated in the rhizosphere when compared to bulk soil transcription (66). In addition to bacteria-plant pathogens, we noted a possible plant-eukaryote interaction. We identified a labelled *Fusarium sp*. (SM_10888_Uncultured_Fungi) in the rhizosphere samples which likely obtained plant-derived carbon directly from the plant through infection.

It appears that not all bacteria growing on root-derived C have identifiable genes that predict a close relationship with the plant. The Stenotrophomonas_67_7, Massilia_66_12, and Burkholderiales_70_21 bins were some of the most highly labelled bins, despite encoding few identifiable interactions systems. Bacteria from these genera are known to be fast-growing soil and rhizosphere associated organisms (67–70). Fast growing organisms may be well positioned to take advantage of the abundance of resources in the rhizosphere.

In addition to plant interaction systems, we also identified genes and pathways that may mediate connections between bacteria and non-plant soil eukaryotes. Several of the eukaryotes we identified were labelled with plant-derived ^13^C, including two nematodes, a rotifer and a rhizaria. Based on phylogeny, these microeukaryotes likely have a bacterivorous lifestyle, and thus probably consumed rhizosphere bacteria living off plant-derived carbon. Several bacterial genes encoded systems that may act as grazing deterrents, for instance the pathway to produce HCN which acts as a nematicidal agent (71). In addition to a type III secretion system with an unknown target, Variovorax_paradoxus_68_14 encodes genes for production of EPS that may defend against predation by micro-eukaryotes, bacteria and phage, provide desiccation protection, improve plant symbiosis, protect the cells from antibiotics and trap nutrients. Bacteria with genomes that encode a large number of fungal cell wall hydrolysis genes may obtain carbon from fungal necromass or actively antagonize living fungi through digestion of their cell walls (72).

In the bacterial genomes we also identified possible mediators of bacteria-bacteria communication and competition. In a previous publication we documented the complete genome of a Saccharibacteria that stole or scavenged nucleotides from rhizosphere bacteria that consumed plant-derived carbon and thus became labelled themselves (38). In labelled genomes we identified many signaling compounds and quorum quenching genes, although at this time we cannot verify their purpose, though it appears communication is likely critical for life in soil (73). The number and distribution of inter-bacterial killing systems in labelled genomes may indicate active competition for resources or space in the rhizosphere. The BGCs that were near genes for plant cell wall depolymerization could produce bioactive compounds that aid in defeating plant defenses or deter other bacteria that could consume the now soluble pool of sugars or prevent other bacteria from colonizing the open plant wounds that they themselves occupy.

One of the most intriguing genomes we reconstructed is for an Acidobacterium. The Acidobacteria_68_21 genome encodes a flexible metabolism and many defensive capabilities, CRISPR Cas systems, insecticidal proteins, and 40 BGCs. While this genome was not labelled, it was one of the most abundant bacterium in bulk soil and these defense systems may serve to protect this dominant bacterium from grazing or parasitism.

In this study we traced fixed carbon from the plant into bacterial genomes, and finally into phage genomes. This enabled us to identify the most active phage in the rhizosphere. We infer that the complete phage genomes are derived from phage particles or phage in the process of replication because of their circularized genomes, rather than phage integrated into bacterial genomes. Interestingly, the two of the most highly labeled phage were more highly isotopically labeled than their bacterial hosts. It is likely that recently synthesized nucleotide pools are more highly labelled that other cell structural components, and these nucleotides were shunted directly into the replicating phage genomes. The presence of highly labelled phage implies that phage predation may be a major source of bacterial death, and thus nutrient cycling, in the rhizosphere.

By tracing the movement of the carbon from the plant into the soil community we can begin to understand rhizosphere ecology, which in turn informs us about the carbon cycle in soil. The possible interactions that we identify in rhizosphere soil have the ability to impact plant growth and shape the flow and stabilization of carbon in soil. Lysis of bacteria by phage or interbacterial killing systems may release easily metabolized compounds that could be respired and returned to the atmosphere. Bacteria may contribute to soil aggregate stability and carbon stabilization through the production of EPS (74). PGPB could enable the plant to fix more CO_2_, ultimately increasing the amount of carbon introduced into the soil (75). Only through a better understanding of the interdomain interactions occurring in soil can we begin to understand the functioning of soil.

## Conclusions

Genome resolved metagenomics applied in the context of a SIP study generated insights into the possible interactive capabilities encoded by the genomes of bacteria, including the subset of soil bacteria that rapidly grow on plant-derived carbon compounds. We thus identified organisms and mechanisms of potential interaction that provide fertile topics for further exploration and verification. Long term, understanding of soil interaction networks may provide pathways to improve plant primary production and carbon compound sequestration in soil.

## Methods

### Labelling

Growing and labelling procedures have been described previously (38). Briefly, soil was collected from the University of California Hopland Research and Extension Center (Hopland, CA, USA), from a field dominated by *Avena spp*. The soil is a fine-loamy, mixed, active, mesic Typic Haploxeralf (76, 77). Microcosms and plant growth conditions have been documented previously (60). A soil sample was collected after microcosm preparation but before planting to represent our time 0 sample. A single *A. fatua* plant was grown in a labeling chamber maintained at 400 μL/L CO_2_, with native CO_2_ replenished with 99 atom% ^13^CO_2_. After six and nine weeks microcosms were destructively harvested. The rhizosphere sample consisted of soil attached to the root after shaking and the bulk soil was collected from a root exclusion mesh bag (1 µm). Rhizosphere soil was washed off the root and DNA was extracted from 0.5g of soil using a phenol:chloroform extraction (60).

### Density separation

We used a CsCl density gradient centrifugation to separate the DNA based on density. Methods have been described previously (78). Briefly, 5.5 μg of DNA was incorporated into a gradient buffer with a density of 1.735 g/mL. The solution was spun in ultracentrifuge tubes (Beckman Coulter Quick-Seal, 13 × 51 mm) in an Optima L-90K ultracentrifuge (Beckman Coulter, Brea, California, USA) using a VTi65.2 rotor at 44,000 rpm (176,284 RCF_avg_) at 20 °C for 109 h with maximum acceleration and braking of the rotor to maintain the integrity of the density separations. The gradient was then gently separated into ∼32 fractions using a syringe pump delivering light mineral oil. Each fraction (∼144 μL) was measured for density using an AR200 digital refractometer (Reichert Inc., Depew, New York, USA). Then DNA was precipitated and quantified as previously described (78). Fractions were then binned based on density and by comparison between the rhizosphere samples and the associated bulk soil (light = 1.692–1.737 g/mL; middle = 1.738–1.746 g/mL; heavy = 1.747–1.765 g/mL; Supplemental figure 1 and Supplemental table 1).

### Sequencing

Each fraction was sequenced at the UC Davis Genome Center on an Illumina HiSeq 3000 (Illumina Inc., Hayward, California, USA) with paired-end libraries prepared with the Kapa Hyper protocol and a read length of 150 bp.

### Sequence preparation and analysis

Reads were trimmed using Sickle (https://github.com/najoshi/sickle) and BBtools (https://sourceforge.net/projects/bbmap/) was used to remove Illumina adapters and trace contaminants. Each sample was assembled individually using IDBA-UD (-step 20, -maxk 140, - mink 40) (79). Only scaffolds larger than 1000 bp were included in future analyses. Genes were predicted using Prodigal (80). The ORFs were annotated using a combined approach. Sequence similarity searches were performed using USEARCH (81) against UniRef100 (82), Uniprot (83), and the KEGG (84) databases. Additional gene annotations were done using HMMs that were constructed based on KEGG Orthologies. All proteins assigned to a KO were clustered using MCL (85) with inflation parameter (-I) of 1.1, based on global percent identity. Clusters were aligned using MAFFT v7 (86), and HMMs were constructed using the HMMER suite (87). Protein domain level analysis was conducted using InterProScan (88). Carbohydrate active enzymes were identified using dbCAN (89). Secondary metabolite clusters were found using antiSMASH (90). tRNAs were predicted using tRNAScan-SE (91). The 16S and 18S rRNA sequences were found and aligned using ssu_tree.py (https://github.com/christophertbrown/bioscripts27). Eukaryotic 18S rRNA genes were dereplicated and clustered at 98% nucleic acid identity representing a possible species level designation (92, 93), aligned using SSU-ALIGN (94) and trees were generated using RAxML on CIPRES (95, 96). Genomes were binned using a combined approach. We used abawaca (https://github.com/CK7/abawaca), MaxBin2 (97), and MetaBAT (98), the best bins were chosen using DAS Tool (99). Further genome curation and binning was done manually on ggKbase (http://www.ggkbase.berkeley.edu). Bins were dereplicated to species level based on rpS3 and 16S rRNA sequence phylogeny.

The rpS3 genes were identified, dereplicated to species level (99% nucleotide identity), and the longest scaffold was chosen using rpS3_trckr (https://github.com/AJProbst/rpS3_trckr). Each sample was mapped to each scaffold using Bowtie2 (--sensitive and --rfg 200,300) and the reads were filtered for two mismatches and the coverage was calculated using calculate_breadth.py (https://github.com/banfieldlab/mattolm-public-scripts/blob/master/calculate_breadth.py) (100). The coverage values for the rpS3 scaffolds were normalized for total read depth from the corresponding sample. The principal coordinate analysis was conducted in the R programming environment with the vegan package (101, 102). The R script is publicly available (103). RpS3 amino acid sequences were aligned using MAFFT v7.402 (86) with the E-INS-i options on Cipres (96). Trees were generated on Cipres using RaxML with the JTT protein substitution model and figures were generated using iTOL (95, 96, 104).

Phage genomes were found using VirSorter and manually on ggKbase (105). Phage genome completeness was checked by mapping reads, as described above, and visualizing on Geneious (106). Circularized phage genomes will have reads paired across the entire span of the scaffold without repetitive elements on the end of the scaffold which could cause long paired reads instead of a circular sequence. CRISPR spacers were found using CRISPRDetect (107).

### Enrichment and qSIP

The coverage and relative abundance of the soil organisms was calculated based on mapping reads using Bowtie2 (--sensitive and --rfg 200,300) (100). For bacterial bins the reads were mapped to all scaffolds in the bin, for eukaryotes the reads were mapped to the whole scaffold containing the 18S rRNA gene, and complete phage genomes were mapped to. Reads were filtered for two mismatches and the coverage was calculated using calculate_breadth.py (https://github.com/banfieldlab/mattolm-public-scripts/blob/master/calculate_breadth.py). The coverage values were normalized for total read depth from the corresponding sample.

We estimated the atom percent excess (APE) ^13^C enrichment for each taxon following the procedures detailed in Hungate et al. (2015), with the following adjustments for metagenome-assembled genomes instead of 16S rRNA genes. Previous qSIP experiments have relied on amplicon sequencing of the 16SrRNA gene to estimate the relative abundances of bacterial and archaeal taxa. Those relative abundances were then converted to estimates of absolute abundance by multiplying by the total number of 16S rRNA gene copies using the universal 16S rRNA primer for qPCR for each density fraction in each replicate gradient. Here, we used the relative coverage (i.e., coverage normalized for read depth) of metagenome-assembled genomes as a proxy for relative abundance and total DNA concentration in place of total 16S copies in order to calculate a metric of abundance (*y*) for each taxon (*i*) in each density fraction (*k*), of each replicate (*j*) as:

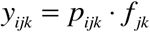

where *p* is relative coverage and *f* is total DNA concentration. This metric of taxon-specific abundance (*y*) is imperfect because total DNA concentration in the rhizosphere presumably represents many more taxa beyond the metagenome-assembled genomes and eukaryotic scaffolds.

Analyses were conducted separately for the week 6 and week 9 data. For each time point, we used the bulk soil data from that week as the unlabeled treatment. This experiment was not replicated, so we were unable to calculate confidence intervals for APE ^13^C, an approach used previously to estimate minimum detectable differences in isotope incorporation (34). However, the relationship between rank and value of APE ^13^C exhibited a spline function (Supplemental figure 7), suggesting a threshold above which label incorporation was detectable above background variation in qSIP-derived estimates of ^13^C uptake. We used breakpoint analysis to test for a spline function, and to identify a threshold value above which ^13^C incorporation could be reasonably inferred. We used the “segmented” package in R, and the segmented function to identify the breakpoint, with the Davies test for significance. We focused this analysis on the 90 taxa exhibiting the lowest values of qSIP-estimated ^13^C atom percent excess (i.e., from ranks 1-90), thereby avoiding more enriched regions where the slope of the rank-APE relationship declined (Supplemental figure 7B). Including the high values skewed the estimate of the breakpoint to near 0 APE, whereas focusing on the data around the visually obvious breakpoint of interest resulted in a more conservative estimate of the threshold. The breakpoint was identified at rank 71.1, with a standard error in this estimate of 0.6. We used the breakpoint plus three times the standard error (the upper 99.7% confidence limit of the breakpoint estimate) to identify the threshold above which ^13^C uptake could be reasonably inferred. The atom percent excess ^13^C value associated with the threshold was 2.5%. Thus, we interpreted values above that APE value as having detectably incorporated the tracer. Note, this is somewhat lower than threshold values estimated using confidence intervals calculated from replicated measurements (5.6% for ^13^C in Hungate et al. AEM).

## Declarations

## Acknowledgements

This work was made possible by samples and expertise provided by personnel at the Hopland Research and Extension Center (Hopland, CA, USA). Plant growth and labeling was conducted at the UC Berkeley Oxford Facility. Labelling, sampling efforts, and technical expertise were provided by Katerina Estera-Molina and Donald J. Herman. Sequencing was done at the UC Davis Genome Center and technical advice was provided by Lutz Froenicke. The HMM pipeline was developed by Dr. David Burstein.

## Contributions

SS and MKF designed the labeling experiment. SS carried out the labeling. ES and SJB conducted the density centrifugation. BJK and BAH conducted the qSIP and growth rate analyses and interpretation. EPS, MKF, and JFB analyzed the data, with input from AJP and JP. EPS, MKF, and JFB wrote the manuscript and all authors read and approved the final manuscript.

## Data availability

The sequences used in this study will be made publicly available upon acceptance of the manuscript.

## Ethics approval and consent to participate

Not applicable.

## Consent for publication

Not applicable.

