## Supplemental Material for "Stable isotope informed genome-resolved metagenomics uncovers potential trophic interactions in rhizosphere soil"

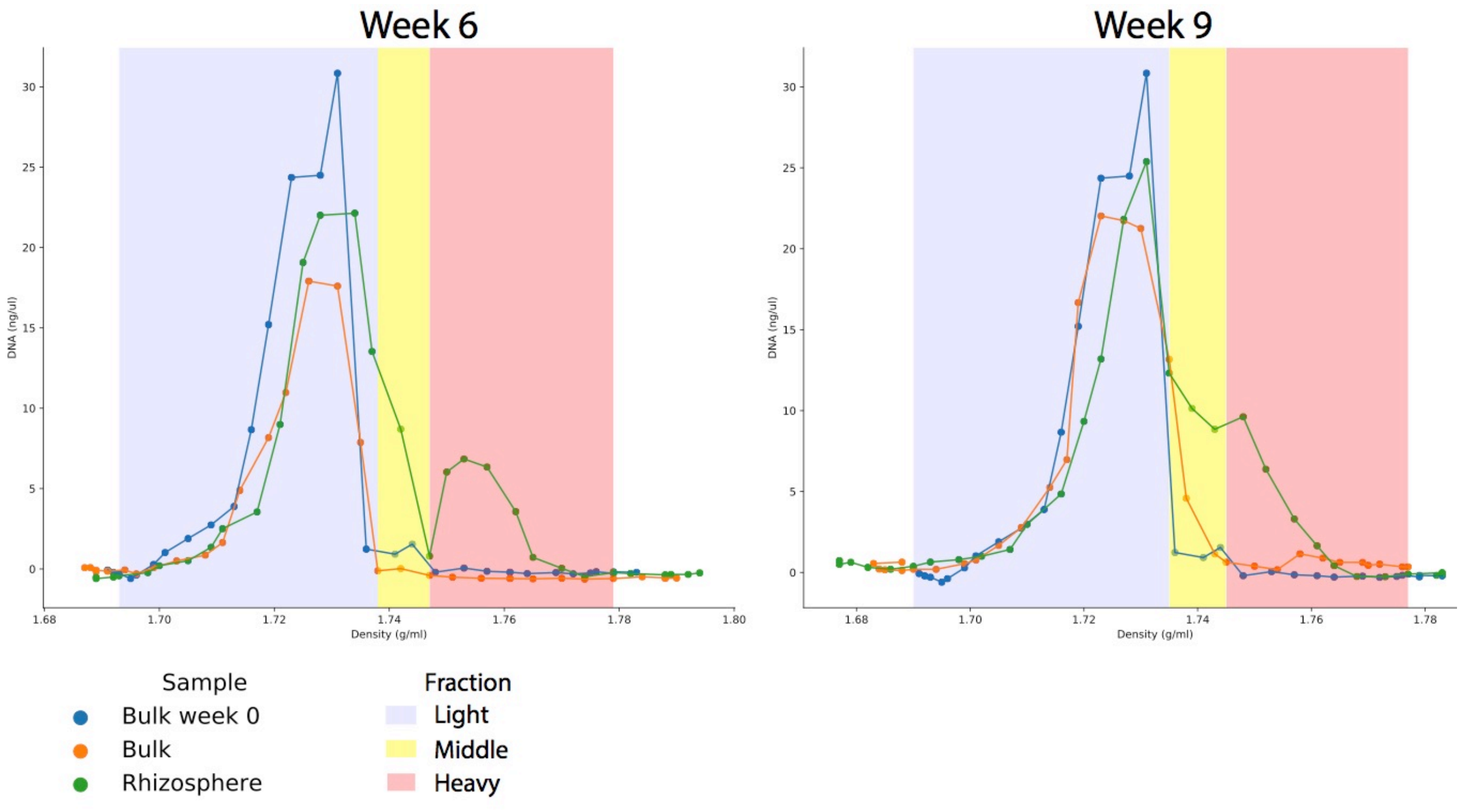


Supplemental figure 1. Stable isotope separation. The density (g/ml) and concentration (ng/µl) of DNA for each fraction collection from each sample is plotted to generate the SIP separation graph.

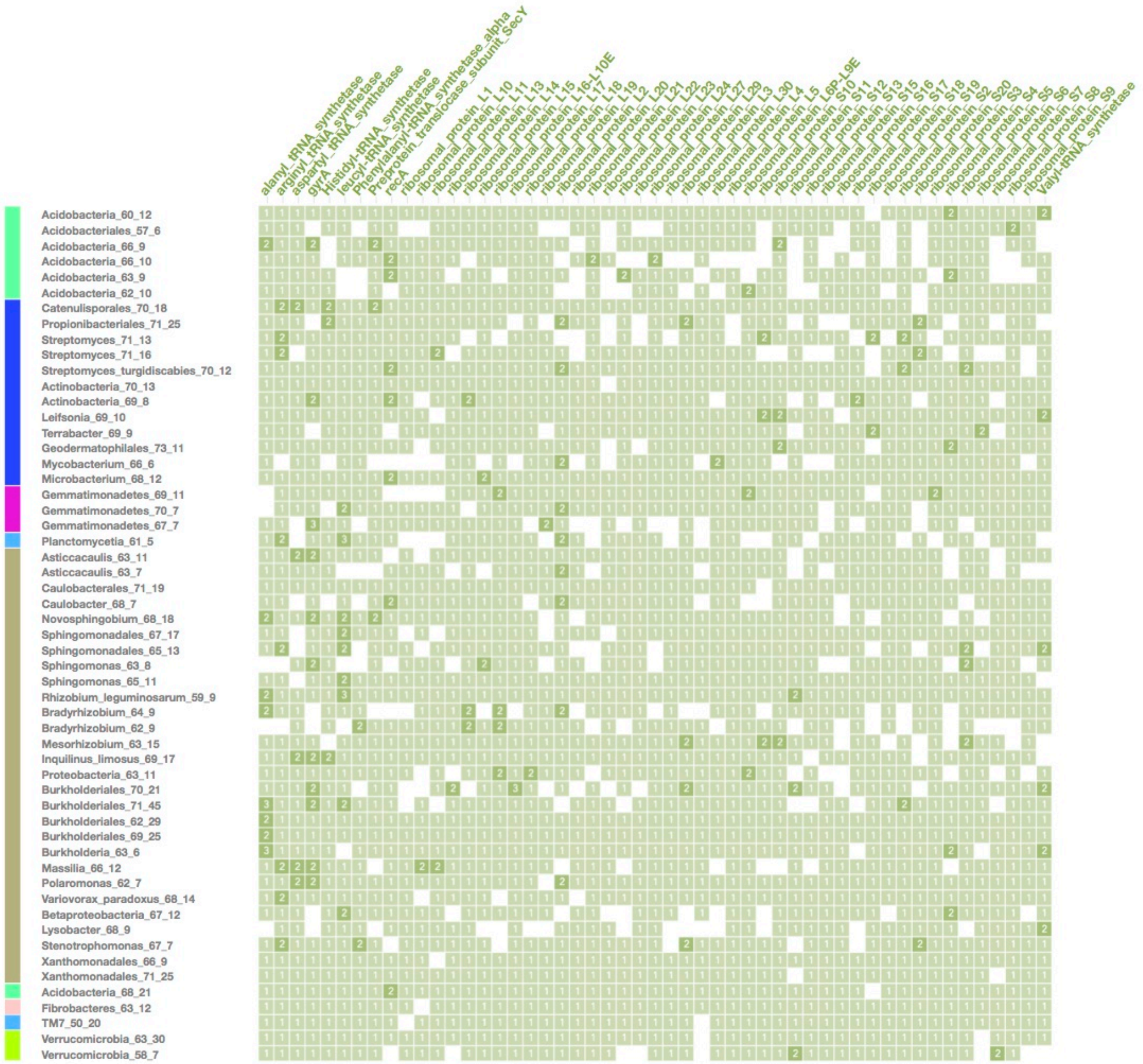


Supplemental figure 2. Count of bacterial single copy genes in the partial bins. Color bar represents clade of bin, colored according to Figure 1.

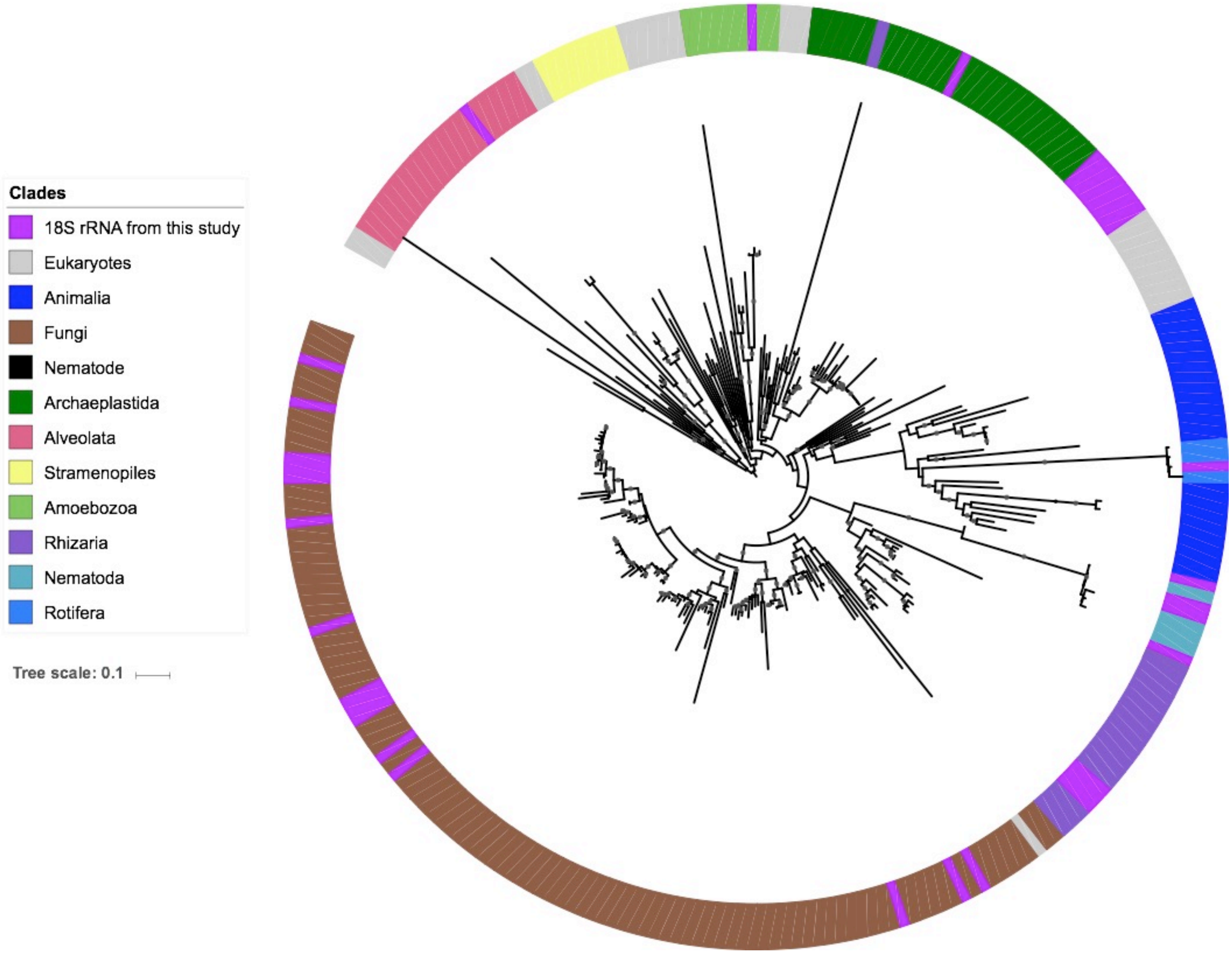


Supplemental figure 3. 18S rRNA tree colored by clade. Black dots represent bootstrap values ≥ 60. Long leaves were removed for ease of visualization and purple indicates 18S rRNA sequences identified in this study

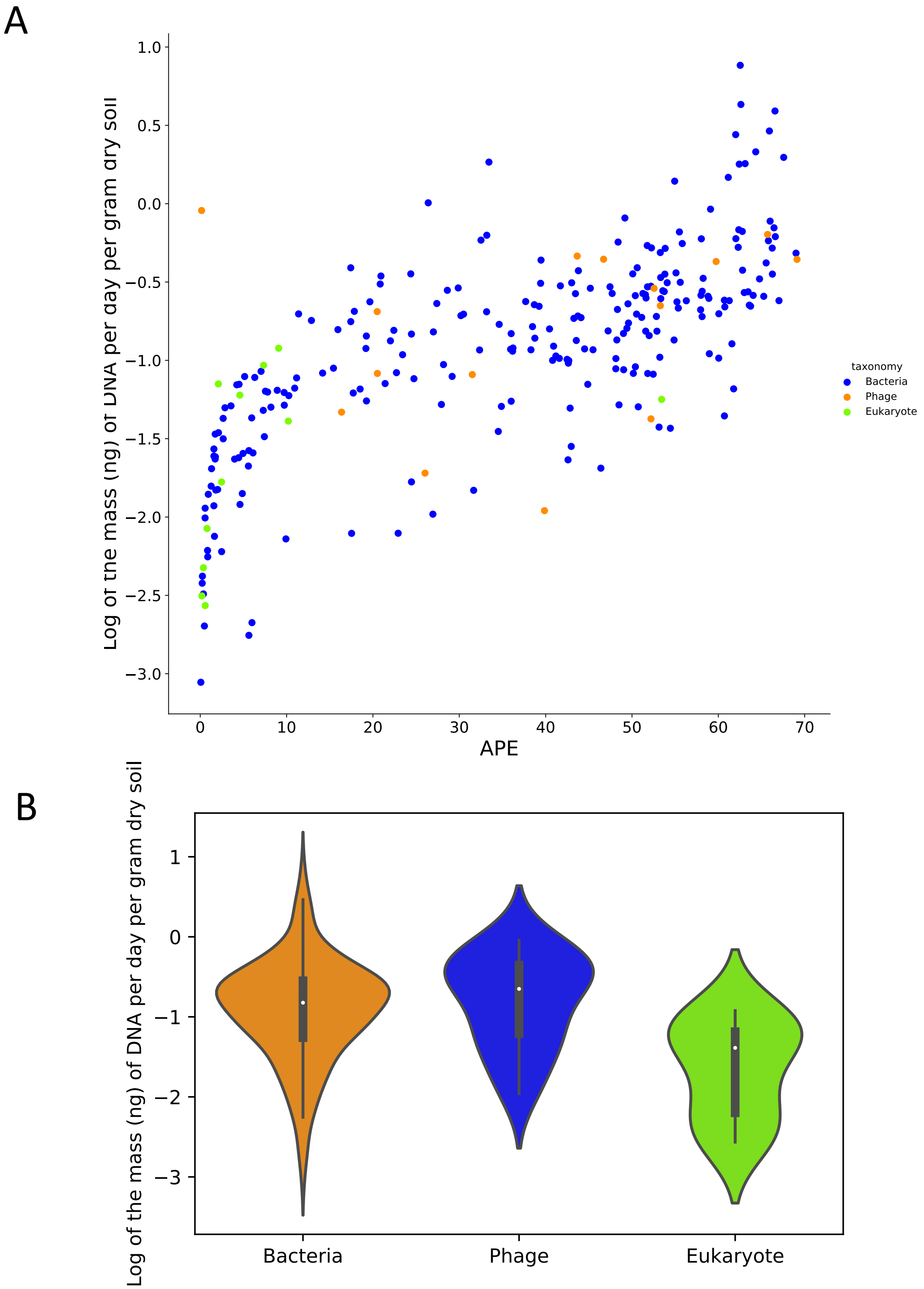


Supplemental figure 4. Gross growth rates of soil organisms on root derived carbon. The log of gross growth per day on plant derived carbon compared to APE (A) and violin plot of gross growth of entities in soil (B).

| Bin | NRPS & NRPS-like | PKS | Bacteriocin | Lanthipeptide | Phosphonate | Indole | Signalling | Terpene | Siderophore | Ectoine | Pigments | Other |
|---|---|---|---|---|---|---|---|---|---|---|---|---|
| Acidobacteria_68_21 | 20 | 3 | 3 | 8 | | | | 1 | | | | 5 |
| Acidobacteria_62_10 | 1 | | | | | | | | | | | |
| Acidobacteria_66_10 | 1 | | | | | | | | | | | |
| Acidobacteria_66_9 | 1 | | | | 1 | | | | | | | |
| Acidobacteriales_57_6 | | | | | | | | | | | | 1 |
| Acidobacteria_60_12 | 1 | 1 | 2 | | | | | 3 | | | | 1 |
| Acidobacteria_63_9 | | | | | | | | 2 | | | | |
| Actinobacteria_69_8 | | | | | | | | | | | | 1 |
| Actinobacteria_70_13 | | | | | | | | 1 | | | | 1 |
| Geodermatophilales_73_11 | 5 | 1 | | | | | | 2 | | | 1 | |
| Propionibacteriales_71_25 | 1 | | | | | | | | | | | |
| Streptomyces_71_16 | 2 | 9 | 2 | | | | | 7 | 2 | 1 | | |
| Catenulisporales_70_18 | 2 | 3 | | 2 | | | | | 1 | | | 2 |
| Terrabacter_69_9 | 1 | 1 | | | | | | | | | | |
| Microbacterium_68_12 | 1 | | | | | | | | 1 | | | 2 |
| Leifsonia_69_10 | 1 | 1 | | | | | | 1 | | | | |
| Streptomyces_71_13 | 11 | 4 | 2 | 2 | | | 3 | 6 | 4 | 1 | 1 | 9 |
| Streptomyces_turgidiscabies_70_12 | 7 | 3 | 2 | 1 | | | | 3 | 2 | 1 | 1 | 2 |
| Mycobacterium_66_6 | 3 | 3 | 1 | | | | | | | | | |
| Caulobacterales_71_19 | | 1 | 1 | | | | | | | | | |
| Caulobacter_68_7 | | | | | | | | | | | | |
| Asticcacaulis_63_11 | | | 2 | | | 1 | | | | | | |
| Asticcacaulis_63_7 | | | 1 | | | 1 | | | | | | 2 |
| Proteobacteria_63_11 | 1 | | | | | | | 1 | | | | |
| Bradyrhizobium_62_9 | | | | | | | | 1 | | | | |
| Bradyrhizobium_64_9 | 5 | | 1 | | | | | 3 | | | | |
| Mesorhizobium_63_15 | | 1 | | | | | 1 | | | | | |
| Rhizobium_leguminosarum_59_9 | | 1 | 2 | | | | 1 | 1 | | 1 | | |
| Sphingomonadales_65_13 | 1 | | | | | | | 2 | | | | 2 |
| Novosphingobium_68_18 | 1 | 1 | | | | | | 1 | | | | 1 |
| Sphingomonadales_67_17 | 2 | | | | | | 2 | 1 | | | | |
| Sphingomonas_65_11 | 1 | 1 | | | | | 1 | 1 | | | | |
| Sphingomonas_63_8 | | | | | | | 1 | | | | | |
| Inquilinus_limosus_69_17 | 3 | 2 | | | 1 | | | 3 | | | | |
| Burkholderia_63_6 | | 1 | 1 | | 1 | | | 3 | | | | |
| Massilia_66_12 | | | 2 | | | | | 1 | 1 | | | |
| Betaproteobacteria_67_12 | 1 | | | | | | | 1 | | | | 1 |
| Variovorax_paradoxus_68_14 | 9 | 1 | 1 | | | | 1 | 1 | | | 2 | 2 |
| Polaromonas_62_7 | | | 1 | | | | | 1 | | | | |
| Burkholderiales_69_25 | 30 | | 2 | | | 1 | 2 | 1 | | | 1 | 3 |
| Burkholderiales_62_29 | 15 | 1 | 2 | | | | 1 | 1 | | | | 2 |
| Burkholderiales_70_21 | 1 | | | | | | 1 | | | | 1 | |
| Burkholderiales_71_45 | | | | | | | | 1 | | | 2 | 1 |
| Xanthomonadales_71_25 | 6 | | 2 | | | | 1 | | | | 1 | 4 |
| Xanthomonadales_66_9 | 1 | | | | | | | | | | 1 | 1 |
| Lysobacter_68_9 | 9 | | 1 | 3 | | | | | | | | |
| Stenotrophomonas_67_7 | 2 | | 2 | | | | | | | | 1 | |
| Gemmatimonadetes_70_7 | | | | | | | | 1 | | | | |
| Gemmatimonadetes_67_7 | | | | | | | | | | | | |
| Gemmatimonadetes_69_11 | 2 | | | | | | | 1 | | | | |
| Planctomycetia_61_5 | | 1 | 1 | | | | | 3 | | | | |
| TM7_50_20 | | | | | | | | | | | | |
| Verrucomicrobia_58_7 | 1 | | | | | | | 3 | | | | |
| Verrucomicrobia_63_30 | | 1 | | | | | | | | | | |
| Fibrobacteres_63_12 | | 1 | 2 | | | | | 1 | | | | |

Supplemental figure 5. Biosynthetic gene clusters encoded in bacterial bins. Signaling compounds includes homoserine lactone clusters, N-acyl amino acid clusters, and butyrolactone cluster.

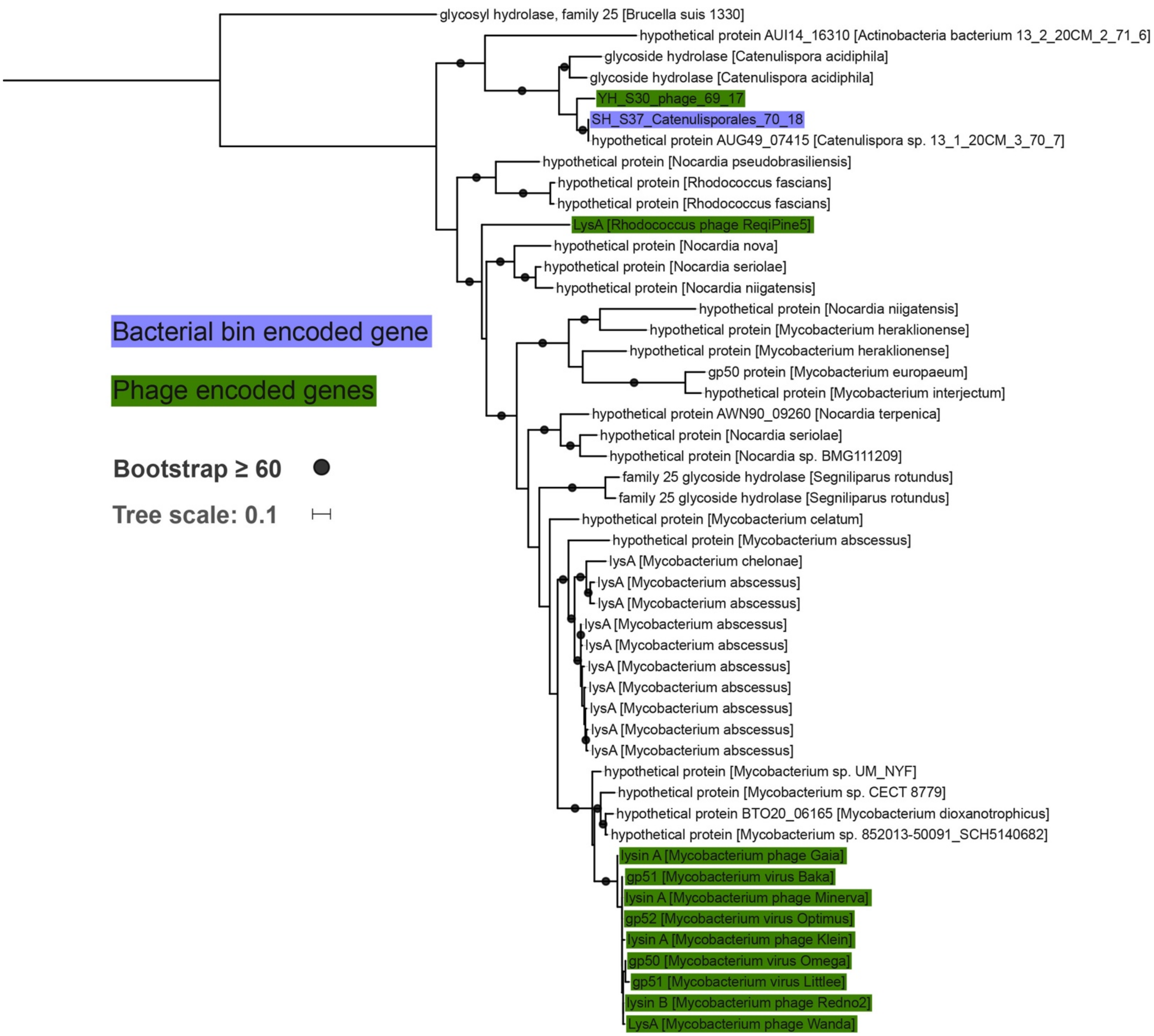


Supplemental figure 6. Phylogenetic tree of GH25 gene. The GH25 genes from phages are highlighted in green and blue highlights the bacterial genome bin from this study.

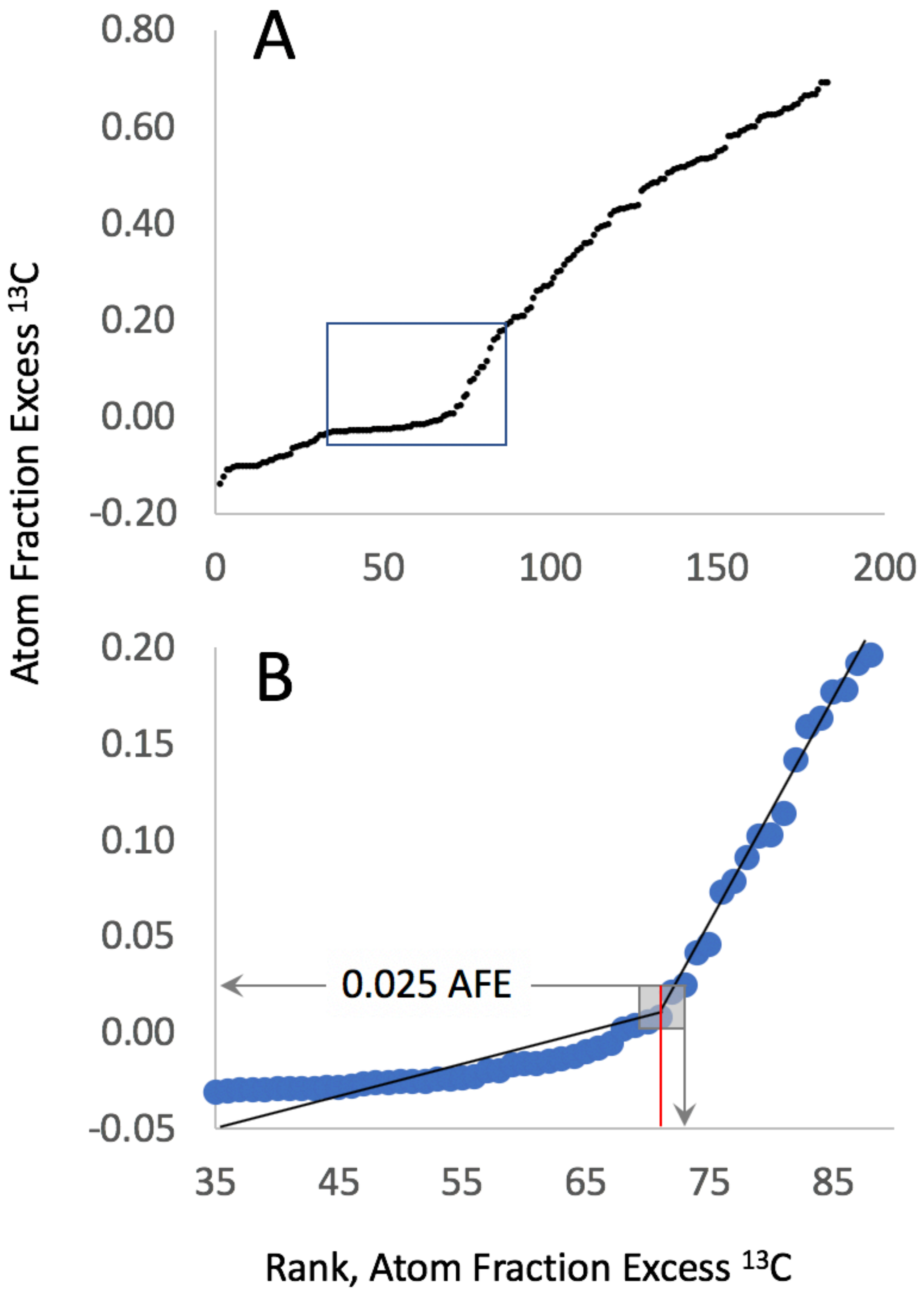


Supplemental figure 7. Break point analysis used to determine enrichment cutoff. Atom faction excess (AFE) compared to rank of organisms based on AFE (A) and inset region (B) focused around the predicted breakpoint region used to calculate the breakpoint for enrichment.

### Supplementary table 1. SIP, sequencing, and assembly statistics.

| Sample | Time point (weeks) | Fraction | Density (g/ml) | DNA concentration (ng/ul) | Total DNA (ng) | Total sequenced (Gbp) | Assembly length (Mbp) | Average contig length (bp) | N50 | Overall alignment rate |
|---|---|---|---|---|---|---|---|---|---|---|
| T0 bulk soil | 0 | Light | 1.695-1.731 | 271 | 2519 | 18.4 | 163 | 1917 | 1922 | 11% |
| T0 bulk soil | 0 | Middle | 1.732-1.744 | 22 | 176 | 16.8 | 267 | 1973 | 1991 | 24% |
| Bulk soil | 6 | Light | 1.694-1.735 | 291 | 2039 | 17.3 | 97 | 1905 | 1801 | 10% |
| Bulk soil | 6 | Middle | 1.736-1.742 | 11 | 109 | 17.4 | 387 | 2209 | 2378 | 30% |
| Rhizosphere soil | 6 | Light | 1.692-1.737 | 267 | 2421 | 16.8 | 112 | 1850 | 1794 | 11% |
| Rhizosphere soil | 6 | Middle | 1.738-1.746 | 17 | 159 | 15.2 | 211 | 2244 | 2385 | 21% |
| Rhizosphere soil | 6 | Heavy | 1.747-1.765 | 137 | 1200 | 17.3 | 470 | 2641 | 3091 | 60% |
| Bulk soil | 9 | Light | 1.694-1.735 | 113 | 2260 | 18.3 | 178 | 2078 | 2078 | 13% |
| Bulk soil | 9 | Middle | 1.736-1.745 | 6 | 127 | 16.4 | 322 | 2129 | 2234 | 26% |
| Rhizosphere soil | 9 | Light | 1.69-1.731 | 82 | 1635 | 18.9 | 176 | 1946 | 1924 | 12% |
| Rhizosphere soil | 9 | Middle | 1.732-1.743 | 31 | 626 | 16.6 | 229 | 2107 | 2108 | 20% |
| Rhizosphere soil | 9 | Heavy | 1.744-1.768 | 21 | 427 | 19.5 | 575 | 2696 | 3314 | 53% |
